# Modelling hCDKL5 heterologous expression in bacteria

**DOI:** 10.1101/2021.06.17.448472

**Authors:** Marco Fondi, Stefano Gonzi, Mikolaj Dziurzynski, Paola Turano, Veronica Ghini, Marzia Calvanese, Andrea Colarusso, Concetta Lauro, Ermenegilda Parrilli, Maria Luisa Tutino

## Abstract

hCDKL5 refers to the human cyclin-dependent kinase that is primarily expressed in the brain where it exerts its function in several neuron districts. Mutations in its coding sequence are often causative of hCDKL5 deficiency disorder. The large-scale recombinant production of hCDKL5 is desirable to boost the translation of current therapeutic approaches into the clinic. However, this is hampered by the following features: i) almost two-thirds of hCDKL5 sequence are predicted to be intrinsically disordered, making this region more susceptible to proteolytic attack; ii) the cytoplasmic accumulation of the enzyme in eukaryotic host cells is associated to toxicity. The bacterium *Pseudoalteromonas haloplanktis* TAC125 (PhTAC125) is the only prokaryotic host in which the full-length production of hCDKL5 has been demonstrated. To date, a system-level understanding of the metabolic burden imposed by hCDKL5 production is missing, although it would be crucial for the upscaling of the production process. Here, we have combined experimental data on protein production and nutrients assimilation with metabolic modelling to infer the global consequences of hCDKL5 production in PhTAC125 and to identify potential overproduction targets. Our analyses showed a remarkable accuracy of the model in simulating the recombinant strain phenotype and also identified priority targets for optimized protein production.

## 1. Introduction

The possibility to heterologously express and purify specific recombinant proteins in large amounts permits their biochemical characterization, the development of commercial goods and their use in industrial processes. With the development of recombinant insulin and its production in *Escherichia coli* in the 1980s [1] a multi-billion dollar market was launched, leading to current large-scale applications that are nowadays capable of releasing products ranging from protein biologics to industrial enzymes [2]. Ideally, the practical steps that lead to recombinant protein production are pretty straightforward and include the identification of the gene of interest, its cloning into an expression vector, its transformation into the host of choice, the induction of protein synthesis and its final purification and characterization [3]. The intrinsic complexity of biological systems, however, usually poses problems down the pipeline of bacterial heterologous protein production. Indeed, as a consequence of the induction of the production of the foreign protein, the biochemistry and physiology of the host may result dramatically altered. The numerous physiological changes that may occur often lowers the amount of the target foreign protein that is produced and eventually recovered from the recombinant organism [4]. In bacteria, high levels recombinant protein production frequently lead to an impact on host cell metabolism; this is usually detectable through growth retardation and is generally known as “metabolic burden” [5]. This additional metabolic load on the microbial chassis has been defined as the portion of a host cell’s resources - either in the form of energy such as ATP or GTP, or raw materials such as amino acids - that is required to maintain and express foreign DNA, as either RNA or protein, in the cell [4]. In *E. coli,* for example, the overexpression of an unnecessary protein results in a linear decrease in the growth rate, with the zero-growth limit occurring when the overexpressed protein occupies a mass fraction equal to 1 - *ϕ_fixed_*, with *ϕ_fixed_* representing the growth-rate invariant fraction of the proteome [6]. There are many factors contributing to the emergence of this burden on growing cells that, at the same time, express a heterologous protein. These mainly include the transcription, the translation and the folding of the foreign protein [5,7,8] and the processes associated with plasmid maintenance, expression and amplification [8,9]. Also, the expression of recombinant proteins may induce a system-level stress response which downregulates key metabolic pathway genes leading to a decline in cellular health and feedback inhibition of both growth and protein expression [10]. Finally, from an energetic perspective, the expression of a foreign protein in a cell may utilize a significant fraction of its metabolic resources and precursors, removing them away from its central metabolism and placing a metabolic drain on the host [4]. Thus, upon protein production induction, an overall cellular reprogramming has to occur in order to ensure an adequate supply of energy and charged amino acids to the process of protein synthesis [10]. The identification of these system-level adjustments following heterologous protein production requires the use of computational representations of microbial metabolism that are able to consider the entire cellular metabolic network. Also, these computational models may help identifying the most suitable approaches to get to target (protein) overproduction. Indeed, it has been recently acknowledged that the most innovative approach currently available to improve the yield of recombinant proteins while minimizing wet lab costs relies on the combination of *in silico* studies to reduce the experimental search space [11]. Among all the available *in silico* approaches, genome-scale metabolic models (GEMs) offers the possibility to predict a cellular phenotype from a genotype under certain environmental conditions and, importantly, to identify possible metabolic targets to improve the production of valuable compounds, while ensuring sufficiently high growth rates [12–14]. GEMs can also be used for descriptive purposes, including the identification of specific metabolic rewiring strategies following external perturbations and/or nutrients switch [15,16]. Thus, not surprisingly, GEMs have been extensively exploited in the context of recombinant protein production, mostly with the aim of optimizing either the cultivation conditions or the strain genetic background for improved recombinant protein production [17–19].

Despite *E. coli* is arguably the bacterium of choice for the production of recombinant proteins, the emergence of novel bacterial chassis is an important fact, especially considering the possible unique properties of their physiology and metabolism, and the practical applications in which they are expected to outperform other microbial platforms [20]. Among them, *Pseudoalteromonas haloplanktis* TAC125 (PhTAC125), the first Antarctic bacterium in which an efficient gene-expression technology was established [21], is particularly promising for a number of reasons. First, several generations of cold-adapted gene-expression vectors allow the production of recombinant proteins either by constitutive or inducible systems and to address the product toward any cell compartment or to the extracellular medium. Secondly, the development of synthetic media and efficient fermentation schemes allows upscaling the recombinant protein production in automatic bioreactors. Finally, the recently reported possibility to produce proteins within a range of temperature from 15 to −2.5 °C enhances the chances to improve the conformational quality and solubility of recombinant proteins. Up to now, PhTAC125 has been used for the production of several recombinant proteins such as a psychrophilic β-galactosidase, the *S. cerevisiae* α-glucosidase, human nerve growth factor and the the lysosomal enzyme a-galactosidase A (hGLA) [22–24].

Recently, PhTAC125 was found to be a potential chassis for the production of human CDKL5 (hCDKL5). hCDKL5 is a cyclin-dependent like protein kinase abundantly expressed in the brain, and it exerts its function in different neuron districts such as the nucleus the cytoplasm and the synaptome. Mutations in the X-linked *cdkl5* gene often end up in the enzyme absence or in the production of loss-of-function variants, and both conditions are causative of hCDKL5 deficiency disorder (CDD), a rare and severe neurodevelopmental disorder for which no cure is available [25]. Recently, a protein replacement therapy was suggested, consisting in the administration of TAT-CDKL5, protein transduction domain (TAT) fused hCDKL5. When injected in *cdkl5* knockout mice, TAT-CDKL5 was able to rescue many anatomical and behavior deficits [26]. The translation of this promising therapeutic approach to clinics needs the large-scale recombinant production of TAT-CDKL5. However, full-length human CDKL5 is a difficult-to-produce enzyme for two main reasons: i) almost two-thirds of its sequence is predicted to be intrinsically disordered, and the lack of a precise 3D structure makes this region more susceptible to proteolytic attack by host-encoded proteases; ii) the cytoplasmic accumulation of the enzyme in eukaryotic cells is associated to considerable toxicity, and the only permissive production strategy is its extracellular secretion, often accompanied with unwanted glycosylation. PhTAC125 is the only prokaryotic cell factory in which the full-length hCDKL5 production has been demonstrated, and the implementation of its efficient production process is the obligatory step towards any possible application (Calvanese et al. 2021 *in press*).

In this work we have modelled the heterologous production of the hCDKL5 protein in the bacterium PhTAC125. The genome-scale model of the recombinant strain was based on its original formulation [27], and further refined/updated and constrained with experimental data on hCDKL5 production and substrates consumption. Such recombinant model was then used to study the global metabolic consequences of the induction of hCDKL5 production as well to identify potential targets for its overproduction.

## 2. Results and Discussion

### 2.1. An updated metabolic reconstruction of PhTAC125

The latest version of ÌMF721 metabolic model of *P. haloplanktis* TAC125 [27] was updated to be compatible with current Systems Biology Markup Language Level 3 Version 2 Core specification [28] extended with Flux Balance Constraints version 2 package specification [29]. The update was conducted with *libsbml* Python library. It covered appropriate objective function declaration, compartments redefinition, model definitions annotation with SBO terms, extension of species definitions with chemical formulas, update of gene names with the newest version of *P. haloplanktis* genome and various minor syntax changes. The update increased iMF721 Memote Total Score from 30% to 78% (Memote reports are available at https://github.com/mdziurzynski/tac125-metabolic-model). Additionally, we used BOFdat [30] to revise the original definition of the biomass composition in iMF721 using available experimental data. More in detail, we used the revised genome sequence of *P. haloplanktis* [31] and a compendium of transcriptomics data from previously published works [32] to improve the formulation of biomass assembly reaction originally proposed [27]. After updating the model, we checked whether it could quantitatively reproduce growth phenotypes as done by the original metabolic reconstruction. Growth simulations on defined media revealed an overall accuracy that matched the one of the original iMF721 reconstruction (Supplementary Material S1). This updated version of the model will be referred to as iMF721_v2 in the following sections and is available at [https://github.com/mfondi/CDKL5_recombinant_production].

### 2.2. CDKL5 production in controlled growth conditions

Human CDKL5 was recombinantly expressed as an N-terminally His-tagged engineered construct to allow for easy Western blot detection and quantification. Its gene was expressed under the control of an IPTG-regulatable promoter [33] cloned in a high-copy number plasmid, named pB40_79C-CDKL5 (average copy number equal to 100, manuscript in preparation) in a mutant version of PhTAC125 - KrPl LacY+ - capable of a fast IPTG internalization [33]. hCDKL5 synthesis was induced in late exponential phase with 5 mM IPTG at 15 °C in bacteria grown in the GG medium [34] for 8 hours. Total production of the target protein was estimated to be 5.2 mg/L of culture by Western blot using a commercial His-tagged calibrator with a similar MW as hCDKL5.

### 2.3. Estimation of average hCDKL5 production flux and nutrients uptake rates

Here, we computed the actual (average) production and growth rates from the experimental data. As for hCDKL5 (molecular weight 128082,77 mg mmol^-1^), after 8 hours a total amount of 5.2 mg (for one liter of culture) was obtained. After the same amount of time, the OD of the culture was measured to be 2.55 that, when multiplied by 0.74 (i.e. the factor for converting PhTAC125 OD to g of biomass [34]) corresponds to 1.887 g of cell dry weight (CDW). Putting everything together, we can compute the average production flux hCDKL5 as follows:

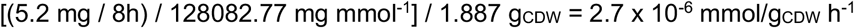

The average growth rate for the recombinant strain across the 8h period was computed using initial and final OD values:

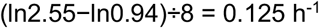

The same calculation led to an average growth rate of 0.169 h^-1^ for the *wt* strain. According to these data, the production of hCDKL5 imposes an overall burden on growing PhTAC125 cells that leads to a 26% reduction in biomass production in the hCDKL5 strain (Figure 1A).

**Figure 1.**
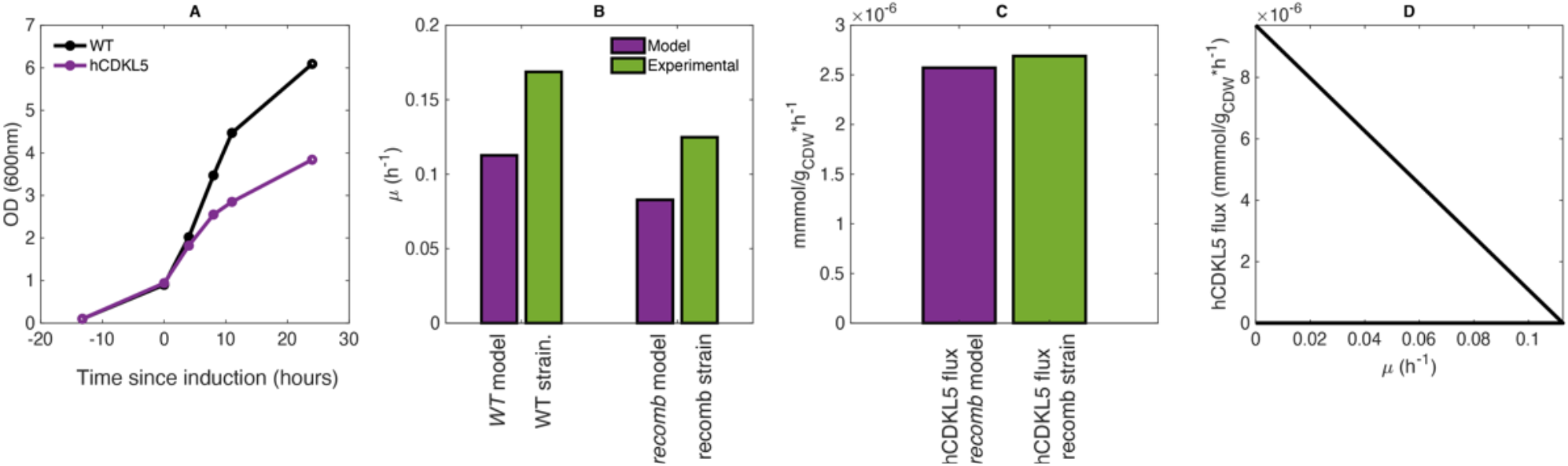
**(a)** Growth curve of WT and hCDKL5 strains as experimentally determined. **(b)** Comparison between the model-predicted and measured growth rates in wild type. **(c)** Comparison between the measured hCDKL5 production rate in the recombinant strain and the one predicted by the model. **(d)** Production enveloper for hCDKL5.

At this point, the only parameters that are missing to fully characterize the CDKL5 production dynamics are the uptake rates for glutamate and gluconate when they represent the only C sources on a minimal medium. To calculate these, we set up an *ad hoc* experiment (see and Material and Methods and Supplementary Material S1) that revealed an uptake rate of 0.35 and 0.66 mmol/g_CDW_*h^-1^ for glutamate and gluconate, respectively.

### 2.4. Recombinant model construction to account for hCDKL5 production

We then extended the iMF721_v2 to include heterologous hCDKL5 production (leading to (resulting in the iMF721_v2_CDKL5 reconstruction, see Supplementary Material S1, Figure S3). The processes taken into account are: (i) synthesis of the plasmid pB40 and (ii) synthesis of hCDKL5 mRNA and its translation into the corresponding protein sequence. As hCDKL5 is not secreted by PhTAC125, no energy-dependent hCDKL5 secretion reaction was added to the model. A plasmid copy number (Pcn) of 100 was used for pB40 because the latter is a high copy-number plasmid. The reaction included in the metabolic network of PhTAC125 representing the synthesis of pB40 is the following:

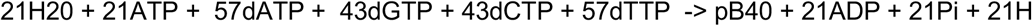

The stoichiometric coefficients for dATP, dGTP, dCTP and dTTP were determined according to the GC composition of the pB40 plasmid. The ATP requirement for the synthesis of the pB40 plasmid was estimated based on the amount of ATP required for the synthesis of the chromosomal DNA as previously described [17,19]. The obtained value (0.21) was multiplied by 100, the estimated copy number of pB40. Finally, pB40 was included in the biomass reaction of the model to account for the burden of the plasmid on the overall physiology of the cell. The stoichiometric coefficient of pB40 was again derived from the stoichiometric coefficient of chromosomal DNA in the biomass assembly reaction of iMF721_v2. This was done using the following proportion: 3850272:0.001608 = 8166:100X, where the first, the second, the third and the fourth terms represent the size (in bp) of the PhTAC125 genome, the stoichiometric coefficient for DNA in the original formulation of the PhTAC125 biomass reaction and the length of the pB40 plasmid, and the (unknown) actual stoichiometric coefficient for the 100 copies of the plasmid. This calculation led to a stoichiometric coefficient for pB40 of 0.000341. Concerning the reaction for hCDKL5 synthesis, this was formalized as follows:

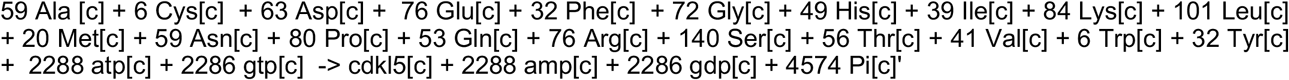

Where the stoichiometric coefficients for the amino acids were based on the composition of protein sequence and the amount of ATP was computed considering the requirement of four ATP molecules for each amino acid added to the protein [35]. As said above, since hCDKL5 is not exported from the cell *in vivo,* no active transport reaction was included in the model.

At this point, we constrained this iMF721_v2_CDKL5 reconstruction with experimental data to build two further models, i.e. a *wt* model and a recombinant model (named *recomb* for brevity). More specifically, we constructed

1. a *wt* model by constraining the iMF721_v2_CDKL5 reconstruction with glutamate/gluconate uptake rates to the values experimentally determined and setting the biomass assembly reaction as the BOF of the model.
2. a *recomb* model by constraining the iMF721_v2_CDKL5 reconstruction with glutamate/gluconate uptake and growth rates to the values experimentally determined and setting the hCDKL5 production reaction as the BOF of the model.

These two models have been used for all the simulations described below. The schematic representation of the computational steps leading to the two models is reported in Supplementary Material S1, Figure S3)

### 2.5. The PhTAC125 recomb model accurately simulates hCDKL5 production

To account for the predictive capability of the PhTAC125 reconstruction in the context of hCDKL5 production, we computed growth and hCDKL5 production rates in the *wt* and *recomb* models. As said above, the *wt* model was obtained by setting the lower bound of glutamate and gluconate uptake reactions to 0.35 and 0.66 mmol/g_CDW_*h^-1^ (respectively) and performing an FBA simulation using biomass production as the objective function. This *wt* model predicted a growth rate of 0.119 h^-1^ which closely resembles the one experimentally measured (Figure 1B, “WT”). Afterwards, to generate the *recomb* model we maintained the same boundaries for the glutamate and gluconate reactions and constrained the growth rate to 74% of the optimal one predicted by the model (74% of 0.119 h^-^) and optimized for hCDKL5 production (Figure 1A). The simulations using this *recomb* model returned a hCDKL5 production flux of 2.67*10^-6^ mmol/g DCW h^-1^ which accurately resembles the one measured experimentally (2.7 x 10^-6^ mmol/g DCW h^-1^) (Figure 1C). A production envelope analysis correctly revealed that hCDKL5 and biomass production competes for a common pool of nutrients and allowed to sketch the current trade-off between these two cellular objectives (Figure 1D). These data indicate that, when constrained with experimental data, the *recomb* model is capable of providing a stoichiometrically reliable representation of hCDKL5 production in PhTAC125.

### 2.6. PhTAC125 metabolic rewiring following hCDKL5 induction

To explore the extent of PhTAC125 metabolic network rewiring upon the induction of hCDKL5 synthesis, we then analyzed the differences in flux distributions between *wt* and *recomb* models. As expected, running an FBA simulations on the two models we found a different number of flux-carrying reactions, with the recomb model showing a higher number of *core* reactions (491 vs 484). However, since an FBA solution may not be unique (i.e. alternative flux distributions may still lead to an equally optimal solution), we used Flux Variability Analysis (FVA) to assess the set of *core* reactions in each of the simulations (see Material and Methods). A set of 84 *core* reactions was shared by the *wt* and the *recomb* models. This set of reactions represents 74% and 97% of the *core* reactions of the two models (i.e. the set of reactions remaining after removing the set of reactions showing a large variability range). Within this set, we identified 12 reactions (11 of them were gene-encoded) shared by both models but that showed an increased flux in the *recomb* vs the *wt* model (Table 1). The 11 gene-encoded reactions included the reactions involved in histidine biosynthesis and an ammonia transporter. The histidine biosynthetic reactions covered the entire pathway, i.e. from PRPP (5-Phosphoribosyl 1-pyrophosphate) to histidine. The higher flux predicted in the histidine biosynthetic pathway of the *recomb* model vs. the *wt* model can be explained by the different amino acid composition of recombinant protein with respect to the native PhTAC125 proteome (Figure 2). Indeed, as the abundance of this amino acid is double in hCDKL5 in respect to the PhTAC125 proteome, precursors used to produce histidine in the *recomb* model will be drained faster than in the *wt* model and fluxes around those precursors are expected to be significantly altered [36].

**Figure 2.**
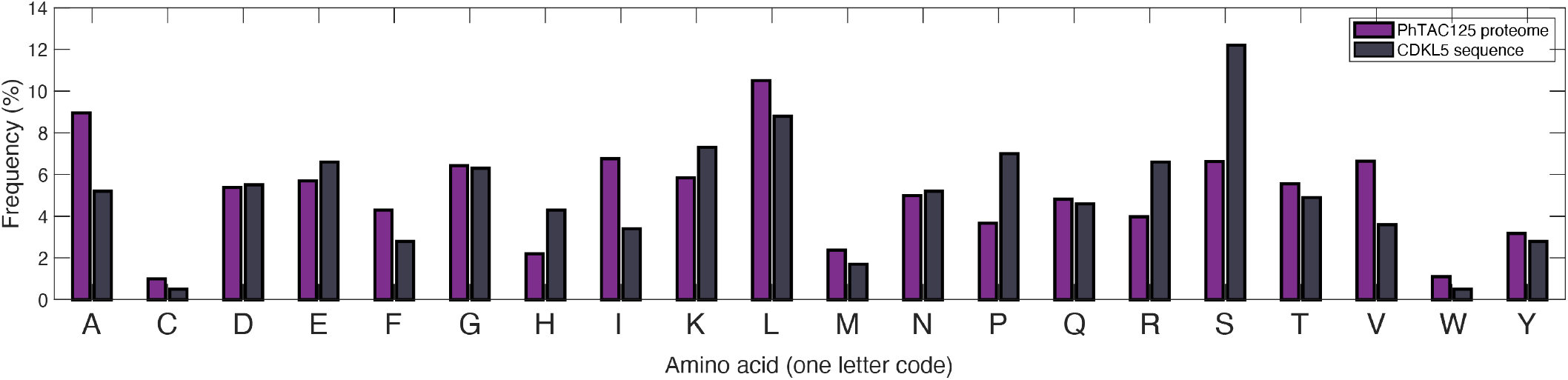
Difference in amino acid composition between PhTAC125 proteome and CDKL5.

**Table 1.** Reactions showing an increased flux in the recomb vs wild model simulations

| Reaction model code | Subsystem | Reaction name |
| --- | --- | --- |
| rxn00789 | Histidine metabolism | 1-(5-Phospho-D-ribosyl)-ATP:pyrophosphate phosphoribosyl-transferase |
| rxn00863 | Histidine metabolism | L-Histidinal:NAD+ oxidoreductase |
| rxn02159 | Histidine metabolism | L-Histidinol:NAD+ oxidoreductase |
| rxn02160 | Histidine metabolism | L-Histidinol-phosphate phosphohydrolase |
| rxn02320 | Histidine metabolism | 5-Amino-2-oxopentanoate:2-oxoglutarate aminotransferase |
| rxn02473 | Histidine metabolism | D-erythro-1-(Imidazol-4-yl)glycerol 3-phosphate hydro-lyase |
| rxn02834 | Histidine metabolism | Phosphoribosyl-ATP pyrophosphohydrolase |
| rxn02835 | Histidine metabolism | 1-(5-phospho-D-ribosyl)-AMP 1,6-hydrolase |
| rxn03135 | Histidine metabolism | Imidazole-glycerol-3-phosphate synthase |
| rxn03175 | Histidine metabolism | N-(5'-Phospho-D-ribosylformimino)-5-amino-1- |
| rxn05466 | Ammonia transport | Ammonia transport via diffusion |

### 2.7. Finding the optimal growth medium

We then sought to identify potential carbon sources whose inclusion in the original, optimized medium, could boost the production of hCDKL5. To this purpose, we selected all the transport reactions present in the iMF721_v2 metabolic reconstruction and created a list including the transported compounds. We considered PhTAC125 as capable of taking up these compounds inside the cell because its genome encodes the corresponding transporters. We then performed one simulation for each of these compounds adding it to the defined medium used during the previous simulations (Schatz salts plus glutamate and gluconate, see Material and Methods), constraining the growth rate to the experimentally determined value and using hCDKL5 production as the objective function. In these simulations the uptake rate of the extra carbon source was arbitrarily set to 0.5 mmol/g_CDW_*h^-1^. We then estimated the effect of the amended carbon source by computing the ratio between the hCDKL5 production flux in the new carbon source and the original one (i.e. with no amendments) and selected the first 30 compounds in the list (Figure 3).

**Figure 3.**
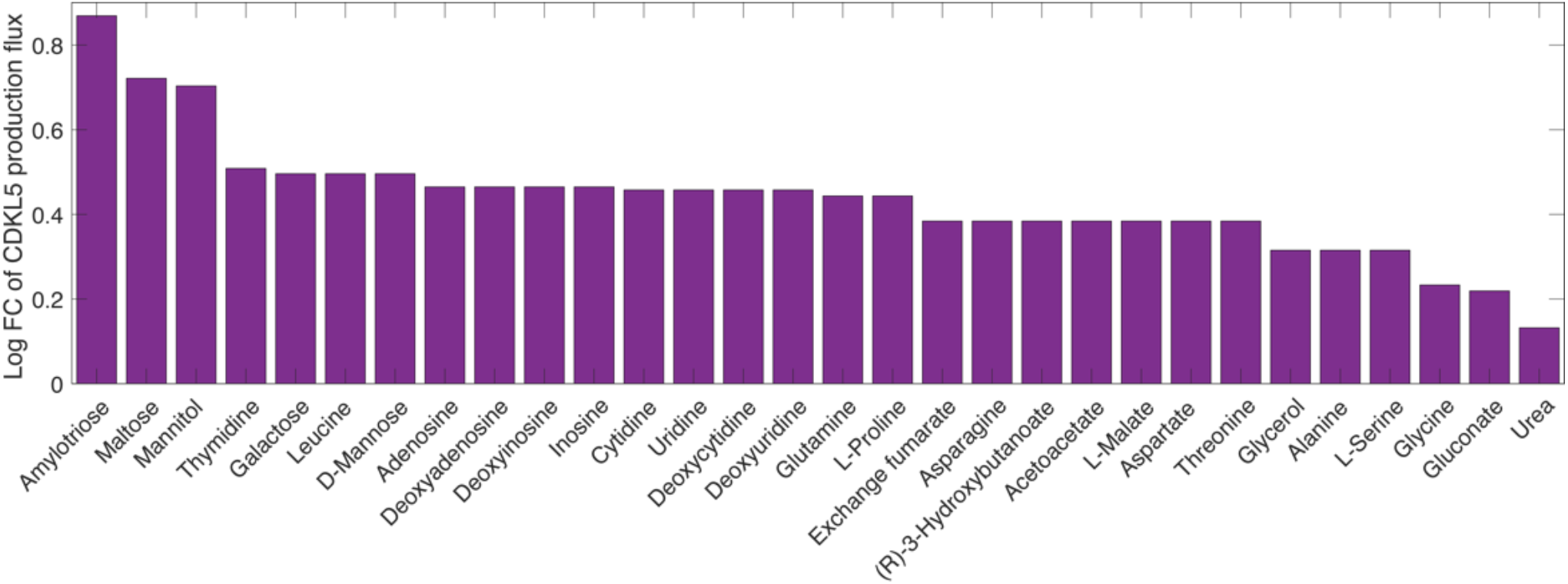
The effect of putative nutrients to be added to the GG medium on CDKL5 production flux. Y-axis indicates the log Fold Change of CDKL5 production flux in repsect to the *recomb* model grown on GG medium.

The three most promising compounds identified with our analysis are amylotriose, maltose and mannitol. The first catabolic steps of these three compounds leads to the formation of D-glucose (amylotriose and maltose) or D-fructose in PhTAC125, thus suggesting that the strengthening of sugar metabolism might the primary effect of adding these compounds to the growth medium of the recombinant strain and one of the possible ways to increase hCDKL5 production. In order to better address this point, we further investigated which part of the PhTAC125 metabolic network is specifically affected by the amendment of the best performing nutrients to the growth medium. We thus checked which reactions increased their flux in the recombinant model growing in GG medium plus amylotriose compared to the same model grown in simple GG medium (Table 2). This list of reactions was filtered by removing those (*non-core*) reactions showing more than 30% variation between their minimum and maximum fluxes during a FVA as described in Material and Methods. Overall, we found 15 gene-encoded reactions displaying an increased flux in this condition: the majority of them (11) were involved in histidine biosynthesis, three in phenylalanine, tyrosine and tryptophan biosynthesis and one in riboflavin metabolism. We found a similar scenario (i.e. the same involved pathways) for the other top four nutrients (maltose, mannitol, thymidine and galactose), with a majority of histidine and phenylalanine metabolism related enzymes displaying an increased flux in the amended medium. Additional pathways that might be affected by these nutrients include nicotinate and nicotinamide metabolism (found when simulating the amendment of thymidine) and galactose metabolism (found when simulating the amendment of galactose).

**Table 2.**
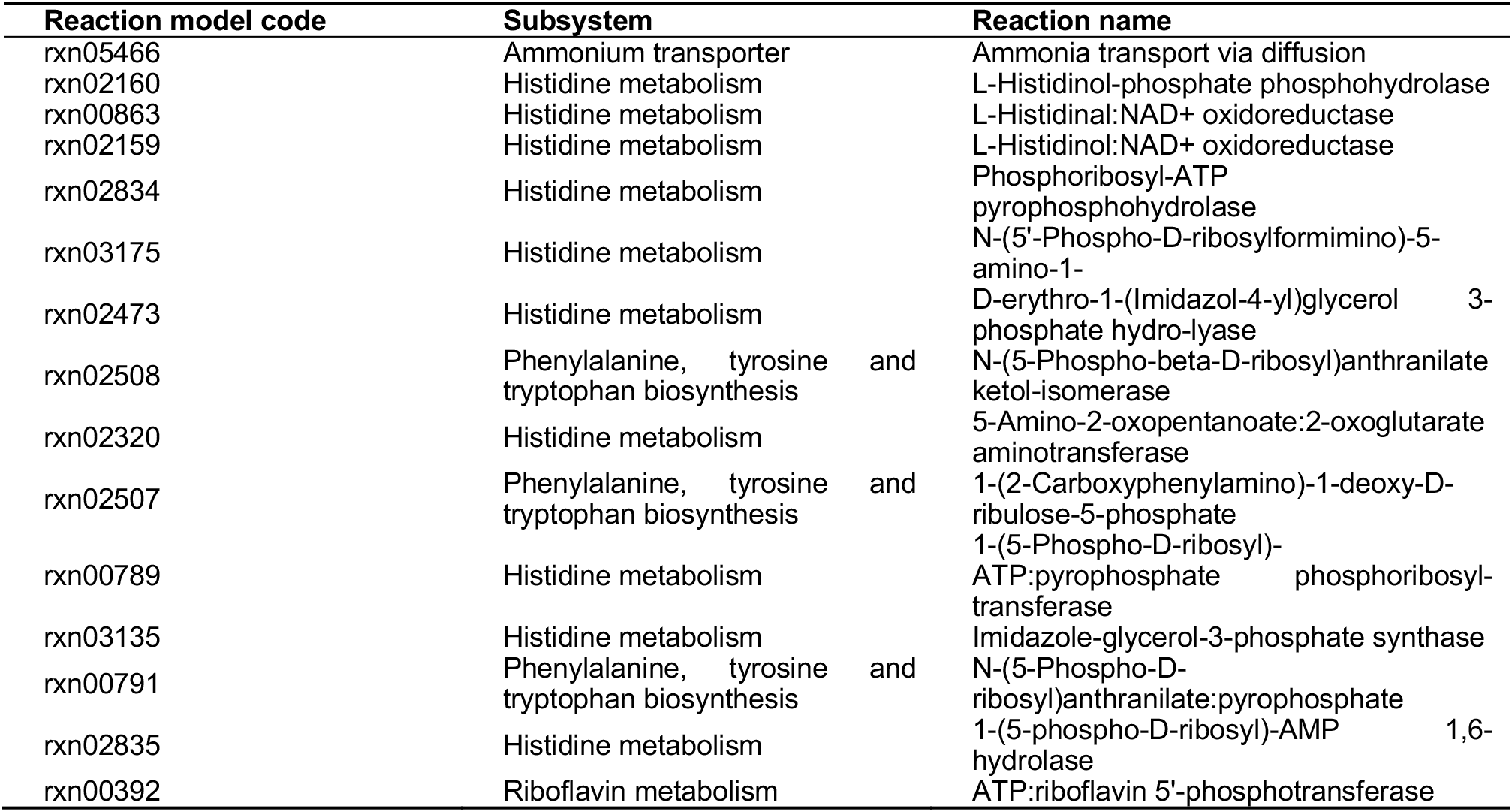
Reactions showing an increased flux in the recomb model growing in GG medium amended with amylotriose

Taken together, these results suggest that the main effect of adding extra-nutrients to the medium would be an increased availability of histidine molecules inside the cell that, in turn, would result in an improved production rate of hCDKL5. Again, this can be explained by the different histidine content of the overall PhTAC125 proteome and of hCDKL5 sequence (Figure 2). The enzymes involved in phenylalanine, tyrosine and tryptophan biosynthesis that appear to increase their flux in the tested conditions include those responsible for the generation of PRPP (5-Phospho-alpha-D-ribose 1-diphosphate) that is a key pentose phosphate pathway (PPP) intermediates for purine, pyrimidine and histidine biosynthesis. Its connection to an increased hCDKL5 production might thus be double: on one side it could fuel histidine biosynthesis for the reasons described above, on the other it could facilitate the synthesis of purines and pyrimidines required by plasmid replication and transcription during heterologous protein expression.

### 2.8. Finding hypothetical targets for hCDKL5 overproduction

We then used the model to predict possible targets to improve the production of hCDKL5 in *P. haloplanktis* TAC125. We focused our attention on the use of the well-established algorithm FSEOF [37]. Briefly, FSEOF scans all the fluxes in the reconstruction and identifies the increasing ones when the flux toward product formation is set (enforced) as further constraint during FBA. The reactions identified by FSEOF are primary over-expression targets that may lead to an improved synthesis of the desired target (hCDKL5 in our case). By applying FSEOF as described in Material and Methods, we identified 70 target gene-encoded reactions, whose over-expression may lead to an improved target production. The complete list of these reactions is available as Supplementary Material S1. The top ten target reactions identified by FSEOF are shown in Table 3. The first reaction in the list is represented by rxn05937, catalyzing the formation of NADPH from NADP and reduced ferredoxin. Forcing the flux through this reaction would allow increasing the overall NADPH pool of the cell and this has been widely recognized as an important factor in the process of heterologous protein production in microorganisms [38]. Reduced ferredoxin necessary for the production of NAPH might be provided by L-Glutamateferredoxin oxidoreductase (Table 3), catalyzing the conversion of L-glutamate to L-glutamine with the reduction of ferredoxin. Reactions belonging to the Entner-Doudoroff (ED) branch of the PPP, are also high-ranking overexpression targets according to FSEOF (Table 3, Figure 4A). These include the three enzymes catalyzing the conversion to D-Glucono-1,5-lactone 6-phosphate to pyruvate and D-Glyceraldehyde 3-phosphate (encoded by *agal, edd* and *eda*). Overall, the degradation of one molecule of glucose through this pathway, as opposed to classical PPP leading to ribose-5P, leads to lower amounts of reducing equivalents (1 NADPH produced instead of 2) but ensure a greater and balanced production of precursors (namely pyruvate and glyceraldehyde-3P (G3P)) that can be used both to fuel the TCA cycle and for amino acid biosynthesis [39]. Indeed, it is known that the ED pathway, as a variant glycolysis pathway, produces equal amounts of G3P and pyruvate and this superior stoichiometric feature makes the ED pathway a preferable route for precursor supply [40]. Importantly, targets within these metabolic pathways (i.e. ED and PPP in general) have been identified in other works aimed at identified optimization production strategies [18,19,39,41]. Most of the other reactions identified by the FSEOF algorithm are involved in the metabolism of amino acids. In particular, our simulations suggest that the production of hCDKL5 might be improved by redirecting the catabolism of glutamate towards the production of aspartate (through the action of L-Aspartate2-oxoglutarate aminotransferase) and its subsequent conversion to 4-Phospho-L-aspartate and L-Aspartate-4-semialdehyde (Figure 4B), catalyzed by ATPL-aspartate 4-phosphotransferase and L-Aspartate-4-semialdehydeNADP+ oxidoreductase, respectively. L-Aspartate-4-semialdehyde, in particular, serves as substrate for the biosynthesis of many amino acids, including lysine, threonine and glycine (Figure 4B). Finally, our FSEOF simulation identified the enzyme serine O-acetyltransferase (catalyzing the formation of serine from CoA and O-Acetyl-L-serine) as a likely hCDKL5 overproduction target. Looking at the unbalanced distribution of S residues in the sequence of hCDKL5 in respect to the one of the PhTAC125 genome (Figure 2) it can be hypothesized that the meaning of this latter finding resides in the necessity to increase the production of serine to cope with the higher request of this amino acid following the induction of CDKL5 production.

**Figure 4.**
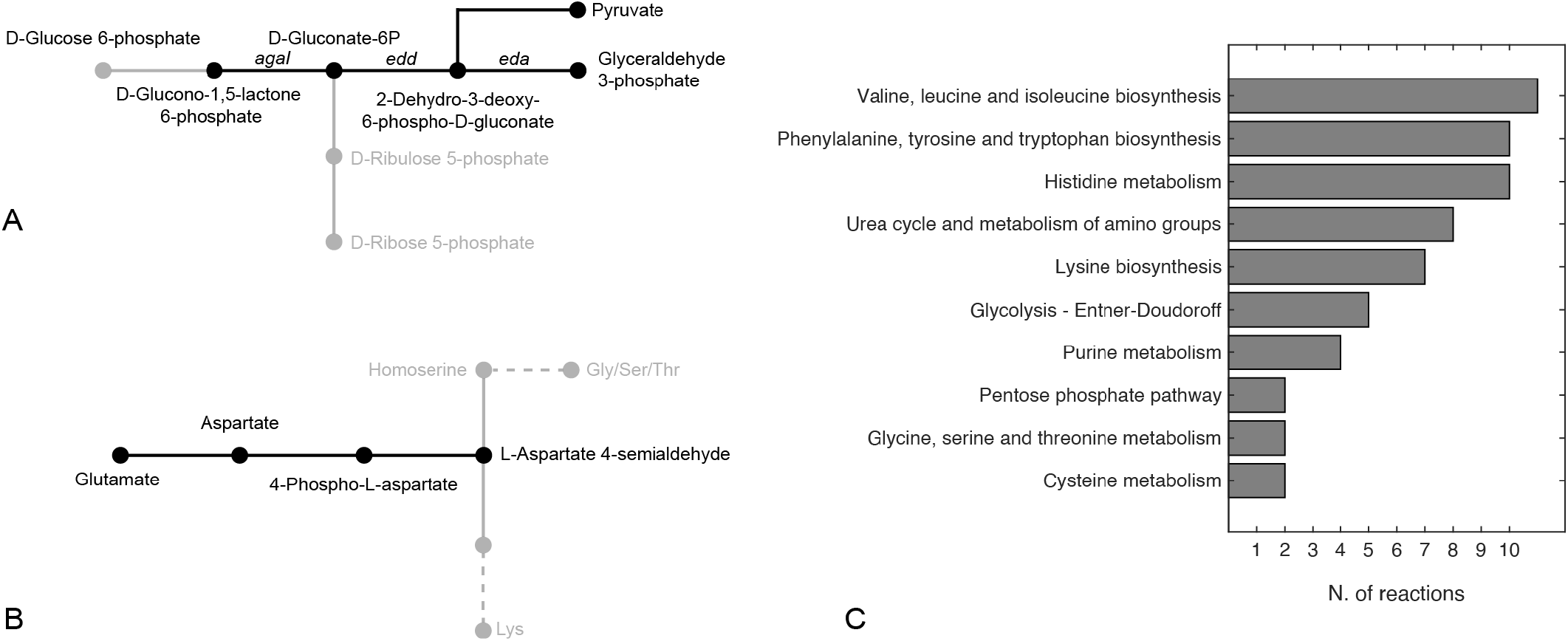
Pathways including the reactions identified as potential over-expression targets by the FSEOF algorithm for **(a)** Entner-Doudoroff pathway and **(b)** glutamate catabolism. Only pathways including 2 or more reactions are shown in **(c)**.

**Table 3.**
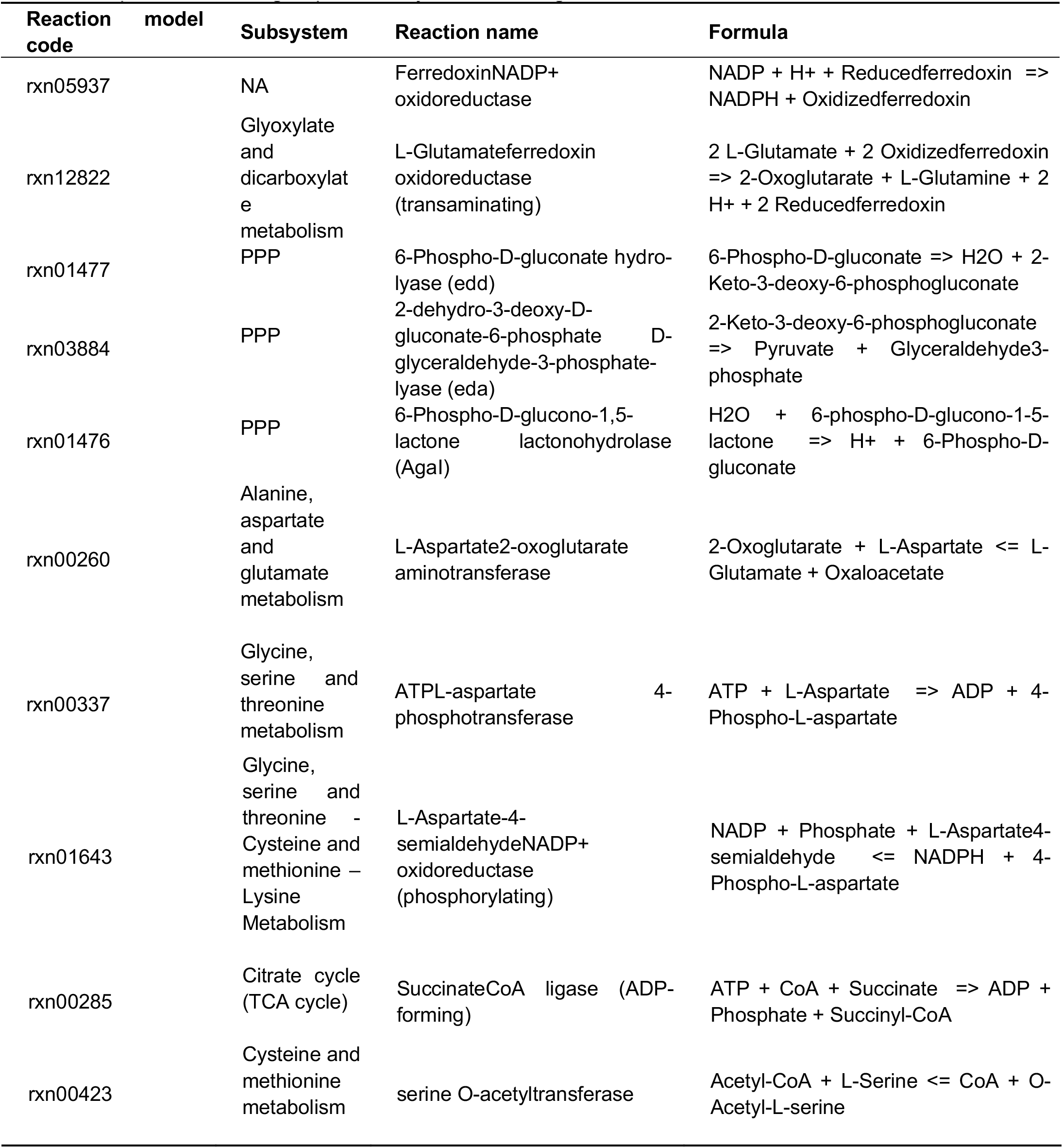
Top ten reaction targets predicted by the FSEOF algorithm

Further, in order to provide a general view of the reactions identified as potential overexpression targets by FSEOF, we grouped them according to their corresponding metabolic pathway (Figure 4C). In line with the results illustrated above, six pathways (out of ten with more than two reactions included) were representatives of amino acids metabolism, with four of them appearing in the top-five pathways (i.e. Val/Leu/Ile, Phe/Tyr/Trp and Lys biosynthesis and His metabolism). Beside amino acid metabolism, the other pathways represented were Urea and amino groups metabolism, glycolysis, PPP and purine metabolism.

## 3. Materials and Methods

### 3.1. Bacterial Strains and conjugation experiments

Plasmid pB40_79C-CDKL5 was mobilized from *E. coli* S17-1(λpir) to KrPL LacY+ [33] through standard conjugation techniques [42]. *E. coli* S17-1(λpir) - a strain possessing *mob* and *tra* genes for plasmid mobilization [43] - was routinely grown in LB (10 g/L bacto-tryptone, 5 g/L yeast extract, 10 g/L NaCl) at 37 °C with the supplementation of 34 μg/mL chloramphenicol if needed for plasmid selection. KrPL LacY+ - a *P. haloplanktis* TAC125 strain engineered for improved IPTG uptake [33] - was grown at 15 °C in TYP (16 g/L bacto-tryptone, 16 g/L yeast extract, 10 g/L NaCl) for conjugational experiments and initial preinocula. Recombinant KrPL LacY+ was selected with 25 μg/mL and 12.5 μg/mL chloramphenicol in liquid and solid media, respectively. Solid LB and TYP broths were prepared by the addition of 15 g/L agar.

### 3.2. hCDKL5 production

pB40_79C-CDKL5 plasmid allows the IPTG-inducible expression of a PhTAC125 codon optimized gene coding for an engineered variant of human CDKL5 isoform 1. The translated protein possesses tandem His-Sumo [44] and Tatk [26,45] N-terminal tags, and a C-terminal 3xflag. The whole 1144 aa sequence was expressed as a cytosolic protein from the pB40 plasmid which is characterized by an average 100 copy number (*manuscript in preparation*). For recombinant gene expression, KrPL LacY+ was cultivated at 15 °C in a 100 mL Erlenmeyer flask containing 20 mL GG medium [34]: 10 g/L L-glutamic acid monosodium salt monohydrate, 10 g/L gluconic acid sodium salt, 10 g/L NaCl, 1 g/L NH4NO3, 1 g/L KH2PO4, 0.2 g/L MgSO4*7H2O, 5 mg/L FeSO4*7H2O, 5 mg/L CaCl2*2H2O, pH 7.8. After the inoculum at 0.10 OD600, the bacterial growth was followed for 13 h and the recombinant gene expression triggered at 1.00 OD600 with 5 mM IPTG. 8 h after the induction, the bacterial cells were harvested by centrifugation (4 °C, 4,000 x*g*, 20 min) when they reached 2.55 OD600. To check and estimate hCDKL5 intracellular production at the end of the culture, bacterial pellets equivalent to 1.00 OD600 were resuspended in 60 μL Laemmli Buffer 4X and denatured at 90 °C for 20 min. Denatured cellular extracts equivalent to 1/120 OD600 were loaded onto a 7.5% precast Mini-Protean TGX (BioRad) and resolved by SDS-PAGE. Known amounts of His-Neuropilin (110 kDa, Immunological Sciences) were loaded onto adjacent lanes to develop a calibration curve. Then, separated proteins were transferred to a PVDF membrane using a semi-dry system and His-tagged proteins (hCDKL5 and His-Neuropilin) were detected with a HRP-conjugated anti-His antibody (1:2,000, Sigma) using the Enhanced Chemiluminescence kit (ECL, BioRad) and a ChemiDoc MP Imaging System (BioRad). Quantitative analyses of blotted hCDKL5 and His-Neuropinilin were carried out with Image Lab software (BioRad) and the volumetric yield was derived considering the final biomass concentration (OD600: 2.55).

### 3.3. Glutamate and gluconate consumption experiment

*Ph*TAC125 bacterial culture was grown in GG medium modified so to contain 5 g/L of L-glutamic acid monosodium salt monohydrate and 5 g/L D-gluconic acid sodium salt in a Stirred Tank Reactor 3 L fermenter (Applikon) with a working volume of 1.5 L. The bioreactor was equipped with the standard pH-, pO2-, level- and temperature sensors for the bioprocess monitoring. The culture was carried out at 15 °C for 30 h in aerobic conditions (45 % dissolved oxygen). 1 mL samples for the metabolomic analysis were collected during the growth and centrifuged at 1300 rpm for 20 min at 4 °C. After the centrifugation, supernatants were recovered, filtered through membranes with a pore diameter of 0.22 μm, and stored at −80 °C.

### 3.4. Metabolomic data

Metabolomic data on cell growth media were obtained by 1H Nuclear Magnetic Resonance Spectroscopy (NMR). The supernatant samples were thawed at room temperature. 540 uL of each sample were added with 60 μL of a potassium phosphate buffer (1.5 M K2HPO4, 100% (v/v) 2H2O, 10 mM sodium trimethylsilyl [2,2,3,3-2H4] propionate (TMSP), pH 7.4). The mixture was then transferred into 5.00 mm NMR tubes for the subsequent analysis.

Spectral acquisition and processing were performed according to standard procedures [46,47]. One-dimensional (1D) 1H NMR spectra were recorded using a Bruker 600 MHz spectrometer (Bruker BioSpin) operating at 600.13 MHz proton Larmor frequency and equipped with a 5 mm PATXI 1H-13C-15N and 2H-decoupling probe including a z axis gradient coil, an automatic tuning and matching and an automatic and refrigerate sample changer (SampleJet). A BTO 2000 thermocouple served for temperature stabilization at the level of ~0.1 K at the sample. Before measurement, samples were kept for 5 min inside the NMR probe head, for temperature equilibration at 300 K. NMR spectra were acquired with water peak suppression using the 1D standard NOESY pulse sequence (128 scans, 65,536 data points, spectral width of 12,019 Hz, acquisition time of 2.7 s, relaxation delay of 4s and mixing time of 0.01 s). The raw data were multiplied by a 0.3 Hz exponential line broadening before applying Fourier transformation. Transformed spectra were automatically corrected for phase and baseline distortions. All the spectra were then calibrated to the reference signal of TMSP at δ 0.00 ppm using TopSpin 3.5 (Bruker BioSpin srl). The signals deriving from glutamate and gluconate were assigned using an internal NMR spectral library of pure organic compounds; matching between the present NMR spectra and the NMR spectral library was performed using AssureNMR software (Bruker BioSpin srl). Their concentrations were calculated by integrating the corresponding signals in defined spectral range, using a homemade R 3.0.2 script.

### 3.5. PhTAC125 genome-scale metabolic reconstruction and constraint-based simulations

The original *P. haloplanktis* TAC125 genome-scale metabolic reconstruction [27] was used as the starting point of the modelling procedures. This metabolic reconstruction was then updated and quality-checked as described above using BOFdat [30] (for the biomass reaction) and Memote [48] (model consistency evaluation). The recently published genome sequence of *P. haloplanktis* TAC125 [31] was fed into BOFdat *DNA.py* script in order to generate the updated stoichiometric coefficients for As, Ts, Cs, and Gs. Similarly, a compendium of expression (RNAseq) data from previously published [32] datasets was fed into BOFdat *RNA.py* code in order to generate revised and experimentally-based stoichiometric coefficients for RNA building blocks.Constraint-based simulation (e.g. FBA) were performed using COBRA Toolbox v3.0 [49] in MATLAB 2020b and using Gurobi as solver. Overexpression targets were identified using the latest FSEOF version implemented in Raven [50] and selecting 100 iterations and a ratio coefficient of optimal target reaction flux of 0.9. The codes used to run all the simulations are available at https://github.com/mfondi/CDKL5_recombinant_production.

### 3.6. Identification of core reactions

Flux Variability Analysis (FVA) was used to assess the relevance of each reaction when simulating growth and hCDKL5 production. The function *fluxVariability* of the COBRA toolbox was used for this purpose. The following procedure was applied (separately) to both the *wt* and the *recomb* models. First, an FBA optimization was run on the model to predict the flux across each reaction. Afterwards, an FVA simulation with exactly the same constraints of the previous FBA simulation was performed and the flux range for each reaction stored. Then, for each of the two models, only those reactions satisfying the following criterion were labelled as *core* reactions:
with *sol_wt/recomb_* > 0

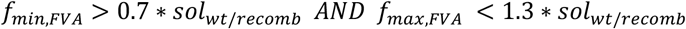

with *sol_wt/recomb_* < 0

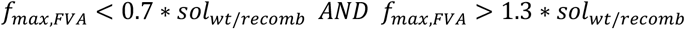

with *sol_wt_*, *f_min,FVA_* and *f_max,FVA_* representing the FBA solution, the lower FVA solution value and the upper FVA solution value, respectively. According to this strategy, in each simulation, only those (*core*) reactions displaying a flux value different from zero and with a narrow range of admissible flux (30%) during a FVA simulation were maintained, whereas those not satisfying this condition were considered unreliable and filtered away.

## 4. Conclusions

In this work, we have combined experimental and computational approaches to characterize the production of recombinant hCDKL5 in the Antarctic marine bacterium *P. haloplanktis* TAC125. By constraining an updated genomescale metabolic model of this bacterium with experimentally determined nutrients absorption rates, we were able to predict hCDKL5 production rates that matched those determined experimentally and to correctly estimate the burden (in terms of reduction in biomass yield, about 25% compared to the *wt* strain) of protein production in this bacterium. Next, we used the model to describe the metabolic rewiring occurring in this bacterium upon the induction of hCDKL5 production and to identify possible overproduction strategies (both in terms of amendments to the original growth medium and in terms of overexpression targets). Despite the fact that each of these analyses highlighted specific pathways and/or targets that appear to be strongly connected to hCDKL5 production, common trends could be identified. In particular, our analyses suggested that the drainage of amino acids and nucleotides towards the synthesis of recombinant hCDKL5 imposes severe alterations in the metabolism of host cell and, as a consequence, the pathways leading to the production of these compounds emerge as priority targets for optimized protein production. In this context, it has been suggested that, when the amino acids composition of the recombinant protein differs from the average composition of the host cell proteome, this might induce an augmented metabolic imbalance in the host cells and an overall bottleneck in the production rate of the recombinant protein [36,51]. In agreement with these observations, we found that i) the metabolism of the *recomb* model is shifted towards histidine biosynthetic fluxes and ii) histidine biosynthetic genes are those that are affected the most by the inclusion of extra nutrients (e.g. amylotriose and maltose) to the medium. Indeed, histidine is one of the residues showing the highest imbalance between the PhTAC125’s proteome and hCDKL5 sequence. Similarly, serine (and other amino acids) biosynthetic steps emerge as priority targets among the reactions identified by FSEOF. Genes and reactions belonging to the PPP are other recurrent hotspots/targets identified by our hCDKL5 production simulations. Studies have previously shown that the overexpression of genes belonging to the PPP have a positive influence on recombinant production in other hosts [39,52] and this has been generally associated to the increased production of both reducing power and intermediates for nucleotides biosynthesis. In our simulations, ED pathway genes (namely *agaI, edd* and *eda*) also emerged as promising targets to increase the rate of hCDKL5 production, probably due to the increased pool of precursors (pyruvate and G3P) available following this route.

## References

1. Goeddel, D.V.; Kleid, D.G.; Bolivar, F.; Heyneker, H.L.; Yansura, D.G.; Crea, R.; Hirose, T.; Kraszewski, A.; Itakura, K.; Riggs, A.D. Expression in Escherichia Coli of Chemically Synthesized Genes for Human Insulin. Proc Natl Acad Sci U S A 1979, 76, 106–110, doi:10.1073/pnas.76.1.106.

2. Burdette, L.A.; Leach, S.A.; Wong, H.T.; Tullman-Ercek, D. Developing Gram-Negative Bacteria for the Secretion of Heterologous Proteins. Microb Cell Fact 2018, 17, 196, doi:10.1186/s12934-018-1041-5.

3. Rosano, G.L.; Ceccarelli, E.A. Recombinant Protein Expression in Escherichia Coli: Advances and Challenges. Front Microbiol 2014, 5, 172, doi:10.3389/fmicb.2014.00172.

4. Glick, B.R. Metabolic Load and Heterologous Gene Expression. Biotechnology Advances 1995, 13, 247–261, doi:10.1016/0734-9750(95)00004-A.

5. Li, Z.; Rinas, U. Recombinant Protein Production Associated Growth Inhibition Results Mainly from Transcription and Not from Translation. Microb Cell Fact 2020, 19, 83, doi:10.1186/s12934-020-01343-y.

6. Scott, M.; Hwa, T. Bacterial Growth Laws and Their Applications. Current Opinion in Biotechnology 2011, 22, 559–565, doi:10.1016/j.copbio.2011.04.014.

7. Bentley, W.E.; Mirjalili, N.; Andersen, D.C.; Davis, R.H.; Kompala, D.S. Plasmid-Encoded Protein: The Principal Factor in the “Metabolic Burden” Associated with Recombinant Bacteria. Biotechnol. Bioeng. 1990, 35, 668–681, doi:10.1002/bit.260350704.

8. Hoffmann, F.; Rinas, U. Stress Induced by Recombinant Protein Production in Escherichia coli. In Physiological Stress Responses in Bioprocesses; Advances in Biochemical Engineering/Biotechnology; Springer Berlin Heidelberg: Berlin, Heidelberg, 2004; Vol. 89, pp. 73–92 ISBN 978-3-540-20311-7.

9. Grabherr, R.; Nilsson, E.; Striedner, G.; Bayer, K. Stabilizing Plasmid Copy Number to Improve Recombinant Protein Production. Biotechnol. Bioeng. 2002, 77, 142–147, doi:10.1002/bit.10104.

10. Sharma, A.K.; Shukla, E.; Janoti, D.S.; Mukherjee, K.J.; Shiloach, J. A Novel Knock out Strategy to Enhance Recombinant Protein Expression in Escherichia Coli. Microb Cell Fact 2020, 19, 148, doi:10.1186/s12934-020-01407-z.

11. Santos, J.; Cardoso, M.; Moreira, I.S.; Gonçalves, J.; Correia, J.D.G.; Verde, S.C.; Melo, R. Integrated in Silico and Experimental Approach towards the Design of a Novel Recombinant Protein Containing an Anti-HER2 ScFv. IJMS 2021, 22, 3547, doi:10.3390/ijms22073547.

12. Fang, X.; Lloyd, C.J.; Palsson, B.O. Reconstructing Organisms in Silico: Genome-Scale Models and Their Emerging Applications. Nat Rev Microbiol 2020, doi:10.1038/s41579-020-00440-4.

13. Zielinski, D.C.; Patel, A.; Palsson, B.O. The Expanding Computational Toolbox for Engineering Microbial Phenotypes at the Genome Scale. Microorganisms 2020, 8, doi:10.3390/microorganisms8122050.

14. Gu, C.; Kim, G.B.; Kim, W.J.; Kim, H.U.; Lee, S.Y. Current Status and Applications of Genome-Scale Metabolic Models. Genome Biol 2019, 20, 121, doi:10.1186/s13059-019-1730-3.

15. Presta, L.; Bosi, E.; Mansouri, L.; Dijkshoorn, L.; Fani, R.; Fondi, M. Constraint-Based Modeling Identifies New Putative Targets to Fight Colistin-Resistant A. Baumannii Infections. Sci Rep 2017, 7, 3706, doi:10.1038/s41598-017-03416-2.

16. Fondi, M.; Bosi, E.; Presta, L.; Natoli, D.; Fani, R. Modelling Microbial Metabolic Rewiring during Growth in a Complex Medium. BMC Genomics 2016, 17, 970, doi:10.1186/s12864-016-3311-0.

17. Sohn, S.B.; Graf, A.B.; Kim, T.Y.; Gasser, B.; Maurer, M.; Ferrer, P.; Mattanovich, D.; Lee, S.Y. Genome-Scale Metabolic Model of Methylotrophic Yeast *Pichia Pastoris* and Its Use for *in Silico* Analysis of Heterologous Protein Production. Biotechnology Journal 2010, 5, 705–715, doi:10.1002/biot.201000078.

18. Nocon, J.; Steiger, M.G.; Pfeffer, M.; Sohn, S.B.; Kim, T.Y.; Maurer, M.; Rußmayer, H.; Pflügl, S.; Ask, M.; Haberhauer-Troyer, C.; et al. Model Based Engineering of Pichia Pastoris Central Metabolism Enhances Recombinant Protein Production. Metab Eng 2014, 24, 129–138, doi:10.1016/j.ymben.2014.05.011.

19. Lule, I.; D’Huys, P.-J.; Van Mellaert, L.; Anné, J.; Bernaerts, K.; Van Impe, J. Metabolic Impact Assessment for Heterologous Protein Production in Streptomyces Lividans Based on Genome-Scale Metabolic Network Modeling. Math Biosci 2013, 246, 113–121, doi:10.1016/j.mbs.2013.08.006.

20. Calero, P.; Nikel, P.I. Chasing Bacterial *Chassis* for Metabolic Engineering: A Perspective Review from Classical to Non-Traditional Microorganisms. Microb. Biotechnol. 2019, 12, 98–124, doi:10.1111/1751-7915.13292.

21. Cusano, A.M.; Parrilli, E.; Marino, G.; Tutino, M.L. A Novel Genetic System for Recombinant Protein Secretion in the Antarctic Pseudoalteromonas Haloplanktis TAC125. Microb Cell Fact 2006, 5, 40, doi:10.1186/1475-2859-5-40.

22. Unzueta, U.; Vázquez, F.; Accardi, G.; Mendoza, R.; Toledo-Rubio, V.; Giuliani, M.; Sannino, F.; Parrilli, E.; Abasolo, I.; Schwartz, S.; et al. Strategies for the Production of Difficult-to-Express Full-Length Eukaryotic Proteins Using Microbial Cell Factories: Production of Human Alpha-Galactosidase A. Appl Microbiol Biotechnol 2015, 99, 5863–5874, doi:10.1007/s00253-014-6328-9.

23. Vigentini, I.; Merico, A.; Tutino, M.L.; Compagno, C.; Marino, G. Optimization of Recombinant Human Nerve Growth Factor Production in the Psychrophilic Pseudoalteromonas Haloplanktis. Journal of Biotechnology 2006, 127, 141–150, doi:10.1016/j.jbiotec.2006.05.019.

24. Papa, R.; Rippa, V.; Sannia, G.; Marino, G.; Duilio, A. An Effective Cold Inducible Expression System Developed in Pseudoalteromonas Haloplanktis TAC125. Journal of Biotechnology 2007, 127, 199–210, doi:10.1016/j.jbiotec.2006.07.003.

25. Katayama, S.; Sueyoshi, N.; Inazu, T.; Kameshita, I. Cyclin-Dependent Kinase-Like 5 (CDKL5): Possible Cellular Signalling Targets and Involvement in CDKL5 Deficiency Disorder. Neural Plasticity 2020, 2020, 1–14, doi:10.1155/2020/6970190.

26. Trazzi, S.; De Franceschi, M.; Fuchs, C.; Bastianini, S.; Viggiano, R.; Lupori, L.; Mazziotti, R.; Medici, G.; Lo Martire, V.; Ren, E.; et al. CDKL5 Protein Substitution Therapy Rescues Neurological Phenotypes of a Mouse Model of CDKL5 Disorder. Human Molecular Genetics 2018, 27, 1572–1592, doi:10.1093/hmg/ddy064.

27. Fondi, M.; Maida, I.; Perrin, E.; Mellera, A.; Mocali, S.; Parrilli, E.; Tutino, M.L.; Liò, P.; Fani, R. Genome-Scale Metabolic Reconstruction and Constraint-Based Modelling of the Antarctic Bacterium *P Seudoalteromonas Haloplanktis* TAC125: Modelling of *P. Haloplanktis* TAC125 Metabolism. Environ Microbiol 2015, 17, 751–766, doi:10.1111/1462-2920.12513.

28. Hucka, M.; Bergmann, F.T.; Chaouiya, C.; Dräger, A.; Hoops, S.; Keating, S.M.; König, M.; Novère, N.L.; Myers, C.J.; Olivier, B.G.; et al. The Systems Biology Markup Language (SBML): Language Specification for Level 3 Version 2 Core Release 2. J Integr Bioinform 2019, 16, doi:10.1515/jib-2019-0021.

29. Olivier, B.G.; Bergmann, F.T. SBML Level 3 Package: Flux Balance Constraints Version 2. J Integr Bioinform 2018, 15, doi:10.1515/jib-2017-0082.

30. Lachance, J.-C.; Lloyd, C.J.; Monk, J.M.; Yang, L.; Sastry, A.V.; Seif, Y.; Palsson, B.O.; Rodrigue, S.; Feist, A.M.; King, Z.A.; et al. BOFdat: Generating Biomass Objective Functions for Genome-Scale Metabolic Models from Experimental Data. PLoS Comput Biol 2019, 15, e1006971, doi:10.1371/journal.pcbi.1006971.

31. Qi, W.; Colarusso, A.; Olombrada, M.; Parrilli, E.; Patrignani, A.; Tutino, M.L.; Toll-Riera, M. New Insights on Pseudoalteromonas Haloplanktis TAC125 Genome Organization and Benchmarks of Genome Assembly Applications Using next and Third Generation Sequencing Technologies. Sci Rep 2019, 9, 16444, doi:10.1038/s41598-019-52832-z.

32. Perrin, E.; Ghini, V.; Giovannini, M.; Di Patti, F.; Cardazzo, B.; Carraro, L.; Fagorzi, C.; Turano, P.; Fani, R.; Fondi, M. Diauxie and Co-Utilization of Carbon Sources Can Coexist during Bacterial Growth in Nutritionally Complex Environments. Nat Commun 2020, 11, 3135, doi:10.1038/s41467-020-16872-8.

33. Colarusso, A.; Lauro, C.; Calvanese, M.; Parrilli, E.; Tutino, M.L. Improvement of Pseudoalteromonas Haloplanktis TAC125 as a Cell Factory: IPTG-Inducible Plasmid Construction and Strain Engineering. Microorganisms 2020, 8, 1466, doi:10.3390/microorganisms8101466.

34. Sannino, F.; Giuliani, M.; Salvatore, U.; Apuzzo, G.A.; de Pascale, D.; Fani, R.; Fondi, M.; Marino, G.; Tutino, M.L.; Parrilli, E. A Novel Synthetic Medium and Expression System for Subzero Growth and Recombinant Protein Production in Pseudoalteromonas Haloplanktis TAC125. Appl Microbiol Biotechnol 2017, 101, 725–734, doi:10.1007/s00253-016-7942-5.

35. Boyle, J. Lehninger Principles of Biochemistry (4th Ed.): Nelson, D., and Cox, M. Biochem. Mol. Biol. Educ. 2005, 33, 74–75, doi:10.1002/bmb.2005.494033010419.

36. Carneiro, S.; Ferreira, E.C.; Rocha, I. Metabolic Responses to Recombinant Bioprocesses in Escherichia Coli. Journal of Biotechnology 2013, 164, 396–408, doi:10.1016/j.jbiotec.2012.08.026.

37. Choi, H.S.; Lee, S.Y.; Kim, T.Y.; Woo, H.M. In Silico Identification of Gene Amplification Targets for Improvement of Lycopene Production. AEM 2010, 76, 3097–3105, doi:10.1128/AEM.00115-10.

38. Driouch, H.; Melzer, G.; Wittmann, C. Integration of in Vivo and in Silico Metabolic Fluxes for Improvement of Recombinant Protein Production. Metabolic Engineering 2012, 14, 47–58, doi:10.1016/j.ymben.2011.11.002.

39. Liu, H.; Sun, Y.; Ramos, K.R.M.; Nisola, G.M.; Valdehuesa, K.N.G.; Lee, W.; Park, S.J.; Chung, W.-J. Combination of Entner-Doudoroff Pathway with MEP Increases Isoprene Production in Engineered Escherichia Coli. PLoS ONE 2013, 8, e83290, doi:10.1371/journal.pone.0083290.

40. Li, C.; Ying, L.-Q.; Zhang, S.-S.; Chen, N.; Liu, W.-F.; Tao, Y. Modification of Targets Related to the Entner-Doudoroff/Pentose Phosphate Pathway Route for Methyl-d-Erythritol 4-Phosphate-Dependent Carotenoid Biosynthesis in Escherichia Coli. Microb Cell Fact 2015, 14, 117, doi:10.1186/s12934-015-0301-x.

41. Aminian-Dehkordi, J.; Mousavi, S.M.; Marashi, S.-A.; Jafari, A.; Mijakovic, I. A Systems-Based Approach for Cyanide Overproduction by Bacillus Megaterium for Gold Bioleaching Enhancement. Front. Bioeng. Biotechnol. 2020, 8, 528, doi:10.3389/fbioe.2020.00528.

42. Tutino, M.; Duilio, A.; Parrilli, E.; Remaut, E.; Sannia, G.; Marino, G. A Novel Replication Element from an Antarctic Plasmid as a Tool for the Expression of Proteins at Low Temperature. Extremophiles 2001, 5, 257–264, doi:10.1007/s007920100203.

43. Tascon, R.I.; Rodriguez-Ferri, E.F.; Gutierrez-Martin, C.B.; Rodriguez-Barbosa, I.; Berche, P.; Vazquez-Boland, J.A. Transposon Mutagenesis in Actinobacillus Pleuropneumoniae with a Tn10 Derivative. Journal of Bacteriology 1993, 175, 5717–5722, doi:10.1128/JB.175.17.5717-5722.1993.

44. Marblestone, J.G. Comparison of SUMO Fusion Technology with Traditional Gene Fusion Systems: Enhanced Expression and Solubility with SUMO. Protein Science 2006, 15, 182–189, doi:10.1110/ps.051812706.

45. Flinterman, M.; Farzaneh, F.; Habib, N.; Malik, F.; Gäken, J.; Tavassoli, M. Delivery of Therapeutic Proteins as Secretable TAT Fusion Products. Molecular Therapy 2009, 17, 334–342, doi:10.1038/mt.2008.256.

46. Takis, P.G.; Ghini, V.; Tenori, L.; Turano, P.; Luchinat, C. Uniqueness of the NMR Approach to Metabolomics. TrAC Trends in Analytical Chemistry 2019, 120, 115300, doi:10.1016/j.trac.2018.10.036.

47. Vignoli, A.; Ghini, V.; Meoni, G.; Licari, C.; Takis, P.G.; Tenori, L.; Turano, P.; Luchinat, C. High-Throughput Metabolomics by 1D NMR. Angew. Chem. Int. Ed. 2019, 58, 968–994, doi:10.1002/anie.201804736.

48. Lieven, C.; Beber, M.E.; Olivier, B.G.; Bergmann, F.T.; Ataman, M.; Babaei, P.; Bartell, J.A.; Blank, L.M.; Chauhan, S.; Correia, K.; et al. MEMOTE for Standardized Genome-Scale Metabolic Model Testing. Nat Biotechnol 2020, 38, 272–276, doi:10.1038/s41587-020-0446-y.

49. Heirendt, L.; Arreckx, S.; Pfau, T.; Mendoza, S.N.; Richelle, A.; Heinken, A.; Haraldsdóttir, H.S.; Wachowiak, J.; Keating, S.M.; Vlasov, V.; et al. Creation and Analysis of Biochemical Constraint-Based Models Using the COBRA Toolbox v.3.0. Nat Protoc 2019, 14, 639–702, doi:10.1038/s41596-018-0098-2.

50. Wang, H.; Marcišauskas, S.; Sánchez, B.J.; Domenzain, I.; Hermansson, D.; Agren, R.; Nielsen, J.; Kerkhoven, E.J. RAVEN 2.0: A Versatile Toolbox for Metabolic Network Reconstruction and a Case Study on Streptomyces Coelicolor. PLoS Comput Biol 2018, 14, e1006541, doi:10.1371/journal.pcbi.1006541.

51. Bonomo, J.; Gill, R.T. Amino Acid Content of Recombinant Proteins Influences the Metabolic Burden Response. Biotechnol. Bioeng. 2005, 90, 116–126, doi:10.1002/bit.20436.

52. Nocon, J.; Steiger, M.; Mairinger, T.; Hohlweg, J.; Rußmayer, H.; Hann, S.; Gasser, B.; Mattanovich, D. Increasing Pentose Phosphate Pathway Flux Enhances Recombinant Protein Production in Pichia Pastoris. Appl Microbiol Biotechnol 2016, 100, 5955–5963, doi:10.1007/s00253-016-7363-5.

