## Supplementary Information S1 for "Modelling hCDKL5 heterologous expression in bacteria"

### Comparison of predicted growth rates with different versions of PhTAC125 genome-scale reconstruction

In order to evaluate whether the changes made to the original formulation of the model, we repeated the simulations reported in Fondi et al. 2014. Results shown below, show that the model is still in good agreement with available experimental data on PhTAC125 growth on minimal media.

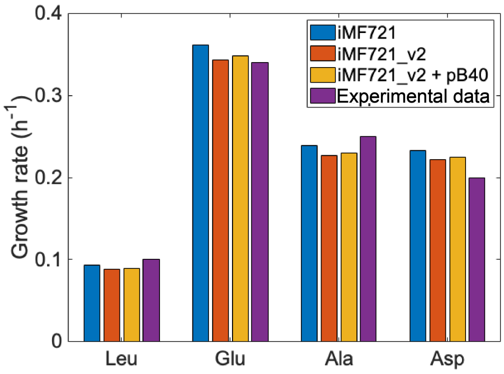

**Figure S1.** Comparison among predicted growth rates of different versions of PhTAC125 metabolic reconstructions. iMF721 refers to the original version of the model as presented in Fondi et al. 2014. iMF721_v2 to the updated version of the model as described in the main text. iMF721_v2 + pB40 refers to this latter model, plus the pB40 plasmid used for CDKL5 protein production.

### Growth and CDKL5 production values

**Table S1.** WT and CDKL5 strains growth parameters over an 8 h growth period.

| | Final OD (8h) | m ($h^{-1}$) |
|---|---|---|
| WT (wt) | 3.47 | 0.169 |
| CDKL5 (recomb) | 2.55 | 0.124 |

### Glutamate and gluconate uptake experimental data

To establish the consumption rate of the carbon sources included in the medium, *Ph*TAC125 bacterial cells were grown GG medium containing 5 g/L of L-glutamic acid monosodium salt monohydrate and 5 g/L D-gluconic acid sodium salt in a Stirred Tank Reactor 3 L fermenter in the conditions described in the main text (Material and Methods). After the growth, the supernatants were recovered, filtered through membranes with a pore diameter of 0.22 μm, and stored at -80 °C before undergoing glutamate and gluconate quantification using NMR as described in the main text. Results of this experiment are reported in Figure S2 together with OD.

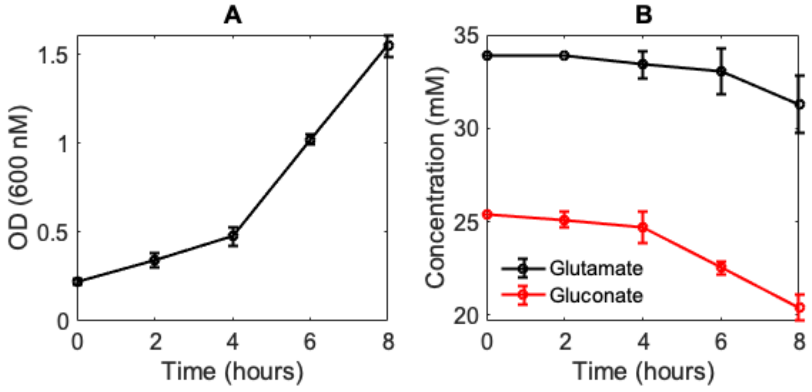

**Figure S2.** Growth (**A**) and glutamate and gluconate consumption (**B**) of PhTAC125 cells over an 8 hours period.

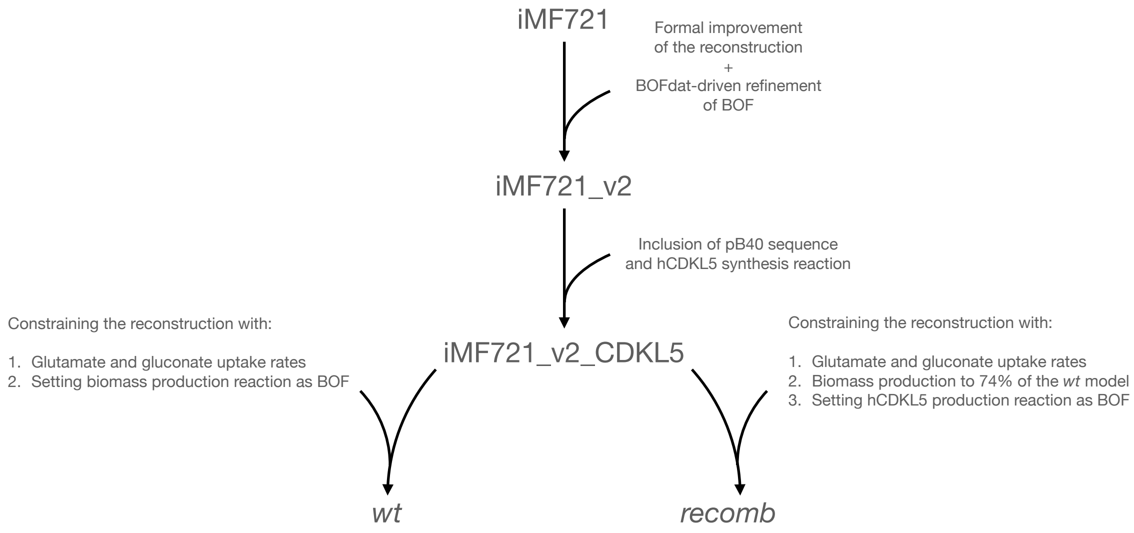

**Figure S3.** The different steps performed in this work to obtain the *wt* and *recomb* metabolic reconstructions.

**Table S2.** Glutamate and gluconate consumption parameters during growth in GG medium with a initial volume of 1.6 L.

| | MW (g/mol) | Initial concentration (g/L) | Final concentration (mM) | Initial concentration (mM) | Yield | Uptake rates (mmol/g DCW $h^{-1}$) |
|---|---|---|---|---|---|---|
| Glutamate | 147.13 | 5 | 31.3 | 33.9 | 0.70 | 0.3463 |
| Gluconate | 196.16 | 5 | 20.4 | 25.4 | 0.36 | 0.6660 |

**Table S3.** The complete list of target reactions identified through FSEOF.

| **Slope** | **rowID** | **Enzyme ID** | **Enzyme Name** | **Direction** | **Gr Rule** |
|---|---|---|---|---|---|
| 4.574 | 7 | rxn00802 | N-(L-Argininosuccinate) arginie-lyase | 1 | PSHA_RS11280 |
| 1.039 | 15 | rxn00011 | pyruvatethiamin diphosphate acetaldehydetransferase | 0 | (( PSHA_RS13615 and PSHA_RS13620 and PSHA_RS01930 ) or ( PSHA_RS17075 and PSHA_RS17070 )) |
| 49.424 | 22 | rxn05466 | Ammonia transport via diffusion | 1 | PSHA_RS11135 |
| 2.416 | 30 | rxn02476 | Phosphoenolpyruvate3-phosphoshikimate | 1 | PSHA_RS07005 |
| 1235,34 | 45 | rxn01985 | Deoxyinosineorthophosphate ribosyltransferase | -1 | PSHA_RS15265 |
| 6.436 | 54 | rxn02186 | 2,3-Dihydroxy-3-methylbutanoateNADP+ oxidoreductase (isomerizing) | 1 | PSHA_RS04145 |
| 2.416 | 57 | rxn02212 | 2-Dehydro-3-deoxy-D-arabino-heptonate 7-phosphate phosphate-lyase | 0 | PSHA_RS13355 |
| 3.899 | 63 | rxn02160 | L-Histidinol-phosphate phosphohydrolase | 0 | PSHA_RS17195 |
| 5.707 | 83 | rxn02789 | 2-Isopropylmalate hydro-lyase | 1 | ( PSHA_RS14190 and PSHA_RS14195 ) |
| 5.707 | 107 | rxn00902 | 3-Carboxy-3-hydroxy-4-methylpentanoate 3-methyl-2-oxobutanoate-lyase | -1 | PSHA_RS14205 |
| 5.535 | 111 | rxn00503 | L-1-Pyrroline-5-carboxylateNAD+ oxidoreductase | -1 | PSHA_RS11170 |
| 3.899 | 129 | rxn00863 | L-HistidinalNAD+ oxidoreductase | 1 | PSHA_RS17205 |
| 7.467 | 148 | rxn00423 | serine O-acetyltransferase | -1 | PSHA_RS03150 |
| 1.039 | 150 | rxn03436 | (S)-2-Aceto-2-hydroxybutanoateNADP+ oxidoreductase (isomerizing) | 1 | PSHA_RS04145 |
| 3.899 | 173 | rxn02159 | L-HistidinolNAD+ oxidoreductase | 1 | PSHA_RS17205 |
| 7.467 | 182 | rxn00649 | O3-Acetyl-L-serine acetate-lyase (adding hydrogen sulfide) | -1 | PSHA_RS09475 |

| | | | | | |
|---|---|---|---|---|---|
| 2.416 | 252 | rxn01332 | PhosphoenolpyruvateD-erythrose-4-phosphate | 0 | PSHA_RS17530 |
| 7.461 | 253 | rxn00566 | L-Cysteine L-homocysteine-lyase (deaminating) | -1 | PSHA_RS09510 |
| 1235,34 | 254 | rxn01858 | Deoxyadenosine aminohydrolase | -1 | PSHA_RS00520 |
| 3.899 | 295 | rxn02834 | Phosphoribosyl-ATP pyrophosphohydrolase | 0 | PSHA_RS17160 |
| 3.899 | 302 | rxn03175 | N-(5'-Phospho-D-ribosylformimino)-5-amino-1- | 1 | PSHA_RS17185 |
| 3.899 | 304 | rxn02473 | D-erythro-1-(Imidazol-4-yl)glycerol 3-phosphate hydro-lyase | 0 | PSHA_RS17195 |
| 4.574 | 317 | rxn01637 | N2-Acetyl-L-ornithine2-oxoglutarate aminotransferase | -1 | ( PSHA_RS00975 or PSHA_RS16905 ) |
| 5.545 | 329 | rxn00313 | meso-2,6-Diaminoheptanedioate carboxy-lyase | 0 | ( PSHA_RS00475 or PSHA_RS00895 ) |
| 10.441 | 376 | rxn00285 | SuccinateCoA ligase (ADP-forming) | 0 | ( PSHA_RS08065 and PSHA_RS08060 ) |
| 4.574 | 395 | rxn00192 | Acetyl-CoAL-glutamate N-acetyltransferase | 0 | PSHA_RS11280 |
| 6.436 | 420 | rxn00898 | 2,3-Dihydroxy-3-methylbutanoate hydro-lyase | 0 | ( PSHA_RS13610 or PSHA_RS15485 ) |
| 2.416 | 421 | rxn01739 | ATPshikimate 3-phosphotransferase | 0 | PSHA_RS13360 |
| 301 | 445 | rxn00414 | Carbon-dioxideL-glutamine amido-ligase (ADP-forming, | 0 | ( PSHA_RS06060 and PSHA_RS06065 ) |
| 16.403 | 448 | rxn00260 | L-Aspartate2-oxoglutarate aminotransferase | -1 | ( PSHA_RS06515 or PSHA_RS04660 or PSHA_RS00305 ) |
| 74.285 | 449 | rxn05937 | FerredoxinNADP+ oxidoreductase | 1 | PSHA_RS15270 |
| 664.470 | 450 | rxn05319 | H2Ot5 | -1 | SPONTANEOUS |
| 1.770 | 456 | rxn00527 | L-Tyrosine2-oxoglutarate aminotransferase | -1 | PSHA_RS17200 |
| 5.194 | 471 | rxn01974 | LL-2,6-Diaminoheptanedioate 2-epimerase | 1 | PSHA_RS00470 |
| 5.194 | 473 | rxn01644 | L-Aspartate-4-semialdehyde hydro-lyase (adding pyruvate and | 0 | ( PSHA_RS16725 or PSHA_RS00905 ) |
| 4.574 | 506 | rxn02465 | N-Acetyl-L-glutamate-5-semialdehydeNADP+ 5-oxidoreductase | -1 | PSHA_RS11300 |
| 37.143 | 519 | rxn12822 | L-Glutamateferredoxin oxidoreductase (transaminating) | 1 | PSHA_RS14360 |
| 1.039 | 522 | rxn03437 | (R)-2,3-Dihydroxy-3-methylpentanoate hydro-lyase | 0 | ( PSHA_RS13610 or PSHA_RS15485 ) |
| 3.899 | 525 | rxn02320 | 5-Amino-2-oxopentanoate2-oxoglutarate aminotransferase | -1 | PSHA_RS17200 |
| 2.416 | 534 | rxn01740 | ShikimateNADP+ 5-oxidoreductase | -1 | PSHA_RS00155 |
| 460 | 576 | rxn05651 | sulfate transport in via proton symport | 1 | PSHA_RS06900 |
| 4.574 | 584 | rxn01434 | L-CitrullineL-aspartate ligase (AMP-forming) | 1 | PSHA_RS11285 |

| | | | | | |
|---|---|---|---|---|---|
| 466 | 586 | rxn00693 | 5-MethyltetrahydrofolateL-homocysteine S-methyltransferase | 1 | ( PSHA_RS10965 and PSHA_RS10970 ) |
| 15.768 | 591 | rxn00337 | ATPL-aspartate 4-phosphotransferase | 0 | ( PSHA_RS13395 or PSHA_RS06160 or PSHA_RS02605 or PSHA_RS11720 ) |
| 2.416 | 616 | rxn02213 | 3-Dehydroquinate hydro-lyase | 1 | PSHA_RS01310 |
| 5.194 | 634 | rxn01973 | N-Succinyl-LL-2,6-diaminoheptanedioate amidohydrolase | 1 | PSHA_RS06550 |
| 460 | 686 | rxn00623 | hydrogen-sulfideNADP+ oxidoreductase | -1 | ( PSHA_RS00790 and PSHA_RS00785 ) |
| 21.970 | 697 | rxn01477 | 6-Phospho-D-gluconate hydro-lyase | 0 | PSHA_RS06715 |
| 5.194 | 706 | rxn03031 | Succinyl-CoA2,3,4,5-tetrahydropyridine-2,6-dicarboxylate | 1 | ( PSHA_RS10075 or PSHA_RS00900 ) |
| 1.039 | 716 | rxn03435 | (R)-2,3-Dihydroxy-3-methylpentanoateNADP+ oxidoreductase | -1 | PSHA_RS04145 |
| 1.334 | 757 | rxn00493 | L-Phenylalanine2-oxoglutarate aminotransferase | -1 | PSHA_RS17200 |
| 5.707 | 759 | rxn01208 | 2-Oxo-4-methyl-3-carboxypentanoate decarboxylation | 0 | SPONTANEOUS |
| 3.899 | 766 | rxn00789 | 1-(5-Phospho-D-ribosyl)-ATPpyrophosphate phosphoribosyl-transferase | -1 | PSHA_RS17210 |
| 460 | 783 | rxn05256 | APSPTi | 0 | PSHA_RS00795 |
| 1.334 | 807 | rxn01000 | Prephenate hydro-lyase (decarboxylating) | 0 | PSHA_RS04630 |
| 15.768 | 816 | rxn01643 | L-Aspartate-4-semialdehydeNADP+ oxidoreductase (phosphorylating) | -1 | ( PSHA_RS10285 or PSHA_RS17375 ) |
| 5.194 | 832 | rxn02929 | 2,3,4,5-TetrahydrodipicolinateNADP+ oxidoreductase | -1 | PSHA_RS06055 |
| 5.707 | 854 | rxn02811 | 3-Isopropylmalate hydro-lyase | -1 | ( PSHA_RS14190 and PSHA_RS14195 ) |
| 4.574 | 861 | rxn01019 | Carbamoyl-phosphateL-ornithine carbamoyltransferase | 1 | PSHA_RS11290 |
| 3.899 | 871 | rxn03135 | Imidazole-glycerol-3-phosphate synthase | 0 | ( PSHA_RS17190 and PSHA_RS17165 ) |
| 4.574 | 872 | rxn00469 | N2-Acetyl-L-ornithine amidohydrolase | 1 | PSHA_RS11305 |
| 5.194 | 900 | rxn03087 | N-Succinyl-L-2,6-diaminoheptanedioate2-ocoglytarate | -1 | ( PSHA_RS00975 or PSHA_RS16905 ) |
| 3.899 | 904 | rxn02835 | 1-(5-phospho-D-ribosyl)-AMP 1,6-hydrolase | 1 | PSHA_RS17160 |
| 4.574 | 923 | rxn01917 | ATPN-acetyl-L-glutamate 5-phosphotransferase | 1 | PSHA_RS11295 |
| 21.970 | 927 | rxn03884 | 2-dehydro-3-deoxy-D-gluconate-6-phosphate | 1 | PSHA_RS06720 |

| | | | | | |
|---|---|---|---|---|---|
| 1235,34 | 946 | rxn00836 | IMPpyrophosphate phosphoribosyltransferase | 1 | PSHA_RS02990 |
| 2.416 | 952 | rxn01255 | 5-O-(1-Carboxyvinyl)-3-phosphoshikimate phosphate-lyase | 0 | PSHA_RS04795 |
| 1.039 | 953 | rxn03194 | (S)-2-Aceto-2-hydroxybutanoate pyruvate-lyase (carboxylating) | 1 | (( PSHA_RS13615 and PSHA_RS13620 ) or ( PSHA_RS17075 and PSHA_RS17070 )) |
| 21.970 | 1017 | rxn01476 | 6-Phospho-D-glucono-1,5-lactone lactonohydrolase | 1 | PSHA_RS05660 |
| 216.684 | 1021 | rxn05312 | PIt6 | 1 | PSHA_RS01555 |
| 1.770 | 1025 | rxn01268 | PrephenateNAD+ oxidoreductase(decarboxylating) | 0 | PSHA_RS04625 |
| 460 | 1032 | rxn09240 | Sulfate adenyltransferase | 1 | ( PSHA_RS01040 and PSHA_RS01035 ) |
| 5.707 | 1084 | rxn03062 | 3-IsopropylmalateNAD+ oxidoreductase | 1 | AUTOCOMPLETION |
| 3.104 | 1086 | rxn01256 | Chorismate pyruvatemutase | 1 | AUTOCOMPLETION |
| 538.953 | 1099 | EX_cpd00067_e | EX_H+_e | -1 | |
| 664.470 | 1113 | EX_cpd00001_e | EX_H2O_e | 0 | |
| 460 | 1130 | EX_cpd00048_e | EX_Sulfate_e | -1 | |
| 216.684 | 1140 | EX_cpd00009_e | EX_Phosphate_e | -1 | |
| 49.424 | 1142 | EX_cpd00013_e | EX_NH3_e | -1 | |
| 6.436 | 1267 | rxn00003 | 2-Acetolactate pyruvate-lyase (carboxylating) | -1 | ( PSHA_RS17075 and PSHA_RS17070 ) |
| 100 | 1328 | CDKL5,c | CDKL5,c | 0 | |
| 1 | 1329 | EX_cdkl5[c] | EX_cdkl5[c] | 0 | |
